# Machine learning reveals conserved chromatin patterns determining meiotic recombination sites in plants

**DOI:** 10.1101/2022.07.11.499557

**Authors:** Minghui Wang, Shay Shilo, Adele Zhou, Mateusz Zelkowski, Mischa A. Olson, Ido Azuri, Nurit Shoshani-Hechel, Cathy Melamed-Bessudo, Alexandre P. Marand, Jiming Jiang, James C. Schnable, Charles J. Underwood, Ian R. Henderson, Qi Sun, Jaroslaw Pillardy, Penny M.A. Kianian, Shahryar F. Kianian, Changbin Chen, Avraham A. Levy, Wojciech P. Pawlowski

**Affiliations:** School of Integrative Plant Science, Cornell University, Ithaca, NY 14853, USA; Bioinformatics Facility, Cornell University, Ithaca, NY 14853, USA; Plant and Environmental Sciences Department, The Weizmann Institute of Science, Rehovot 76100, Israel; Department of Plant Biology, Michigan State University, East Lansing, MI 48824, USA; Department of Agronomy and Horticulture, University of Nebraska, Lincoln, NE 68588, USA; Department of Chromosome Biology, Max Planck Institute for Plant Breeding Research, 50829 Cologne, Germany; Department of Plant Sciences, University of Cambridge, Cambridge CB2 3EA, UK; Department of Horticultural Science, University of Minnesota, St. Paul, MN 55108, USA; USDA-ARS, Cereal Disease Laboratory, St. Paul, MN 55108, USA; School of Life Sciences, Arizona State University, Tempe, AZ 85287, USA

## Abstract

Distribution of meiotic recombination events in plants has been associated with local chromatin and DNA characteristics, chromosome landmark proximity, and other features^1-7^. However, relative importance of these characteristics is unclear and it is unknown if they are sufficient to unambiguously determine recombination landscape^8^. Here, we analyzed over 40 DNA sequence, chromatin, and chromosome location features of maize and *Arabidopsis* recombination sites using machine learning^9,10^. We discovered that a combination of just three features, CG methylation, CHG methylation, and nucleosome occupancy, enabled identification of exact crossover site with 90% accuracy. These results imply redundancy of most recombination site characteristics. Recombination takes place in a small fraction of the genome with chromatin features distinct from those of genome at large. Surprisingly, crossover sites show elevated heterochromatin histone marks despite low DNA methylation. Crossover site features show broad evolutionary conservation, which will enable creating genetic maps in species where conventional mapping is unfeasible.

## INTRODUCTION

Meiotic recombination is initiated by the formation of double-strand breaks (DSBs) in chromosomal DNA^11^. DSB repair produces crossovers (COs)^12,13^ by first generating recombination intermediates called double Holliday junctions (dHJ)^14^ that create connections between homologous chromosomes. However, in most species, DSBs vastly outnumber COs and the vast majority of DSBs are repaired as non-COs. In maize, there are ∼ 500 DSBs but fewer than 20 COs per meiosis^15^. Furthermore, not all genomic sites are equally amenable to recombination. What constitutes recombination-competent chromatin exhibits substantial inter-species variation^8^. In mammals, recombination sites are strongly associated with elevated presence of H3K4 tri-methylation (H3K4me3), which results from the action of the PRDM9 protein^16^. In contrast, in plants, which lack PRDM9, the relationship between H3K4me3 and recombination is more nuanced ^1-3,17^.

## RESULTS AND DISCUSSION

To better understand how local chromosome characteristics affect the position of recombination events, we used a set of local DNA sequence, chromatin, and chromosome location features (Table 1, Supplemental Table 1, Supplemental Figures 1 and 2) to dissect high-resolution datasets of DSBs and COs, from maize and Arabidopsis^1-3,17^. The datasets contained ∼3000 DSB hotspots and ∼1100 COs for each species. Characteristics of these sites were compared to those of recombination desert regions. Recombination deserts were defined separately for DSBs and COs. In both cases, we compiled the longest 5% of stretches between adjacent recombination sites, DSBs or COs, respectively. Centromere regions were excluded as they may possess unusual features that block recombination^18^. To avoid bias, the same total DNA length was randomly selected from the desert regions as the total DNA length of the DSB or CO intervals.

**Table 1.** List of genomic and chromatin features associated with recombination sites.

|  |
| --- |
| DNA sequence at recombination site: |
| - SNP density (*for COs only*) |
| - DNA sequence motifs |
| - G/C content |
| - A/T content |
| - dinucleotide repeats (AA, AC, AG, AT, AC, CC, CG, CT, GA, GC, GG, GT, TA, TC, TG, TT) |
| Gene features |
| - overlapping an expressed gene |
| - overlapping a gene expressed in meiosis (*maize only*) |
| - overlapping an exon |
| - overlapping a single-exon gene (*maize only*) |
| - distance to nearest transcription start site (TSS) |
| - distance to nearest transcript termination site (TTS) |
| - overlapping a transposable elements (TE) |
| DNA structure: |
| - minor groove width (MGW) |
| - roll |
| - propeller twist (ProT) |
| - helix twist (HelT) |
| Chromosome features: |
| - distance to telomere |
| - distance to centromere |
| Chromatin features: |
| - H2A.Z (*Arabidopsis only*) |
| - H3K4me3 |
| - H3K9me2 |
| - H3K27me3 |
| - H3K56ac |
| - H4K5ac (*maize only*) |
| - DNA methylation (<sup>m</sup>CG, <sup>m</sup>CHG, <sup>m</sup>CHH-*Arabidopsis only*) |
| - micrococcal nuclease sensitivity (MNase) |
| - Transposase Accessible Chromatin (ATAC) |

Using principle component analysis (PCA), we found that DSB and CO sites in both species showed good separation from recombination desert regions (Supplemental Figure 3). Individual features were also nearly all significantly different between recombination sites and recombination deserts. In maize, of the 42 features examined, only six, CA, CC, CG, TA, and TG dinucleotide repeats, did not show significant differences between recombination sites and recombination deserts (Supplemental Table 2). These data indicate that genomic locations of DSBs and COs exhibit characteristics distinct from those of recombination-poor genome regions.

To determine whether DSB and CO sites can be identified solely based on their features, we examined three ML algorithms, Random Forest, Logistic Regression, and Decision Tree. We evaluated the ability of the algorithms to identify locations of DSB and CO sites with high accuracy, i.e. with the resolution of 2 kb. All three algorithms resulted in high values of Area Under the Curve (AUC), which represents the fraction of correctly-identified recombination sites (Figure 1A). The Random Forest algorithm exhibited the best performance, with AUC ranging from 0.84 to 0.98 (Figure 1A), with 1.0 corresponding to a perfectly-performing algorithm. Algorithm performance was better in maize than in Arabidopsis, possibly because most epigenetic data in maize, in contrast to Arabidopsis, were derived from meiotic tissues. We further tested the Arabidopsis algorithm using a larger CO collection comprised of datasets from Underwood *et al*.^19^ and Rowan *et al*.^20^. While using these datasets for training did not improve ML algorithm performance, using it for cross-validation of the algorithms trained on the original 1,100-CO dataset resulted in equally high accuracy as validation in the original dataset. This result provides additional verification of robustness our ML algorithms. Altogether, our observations indicate that recombination sites can be reliably identified with high precision based on local chromatin, DNA sequence, and chromosome location features.

**Figure 1.**
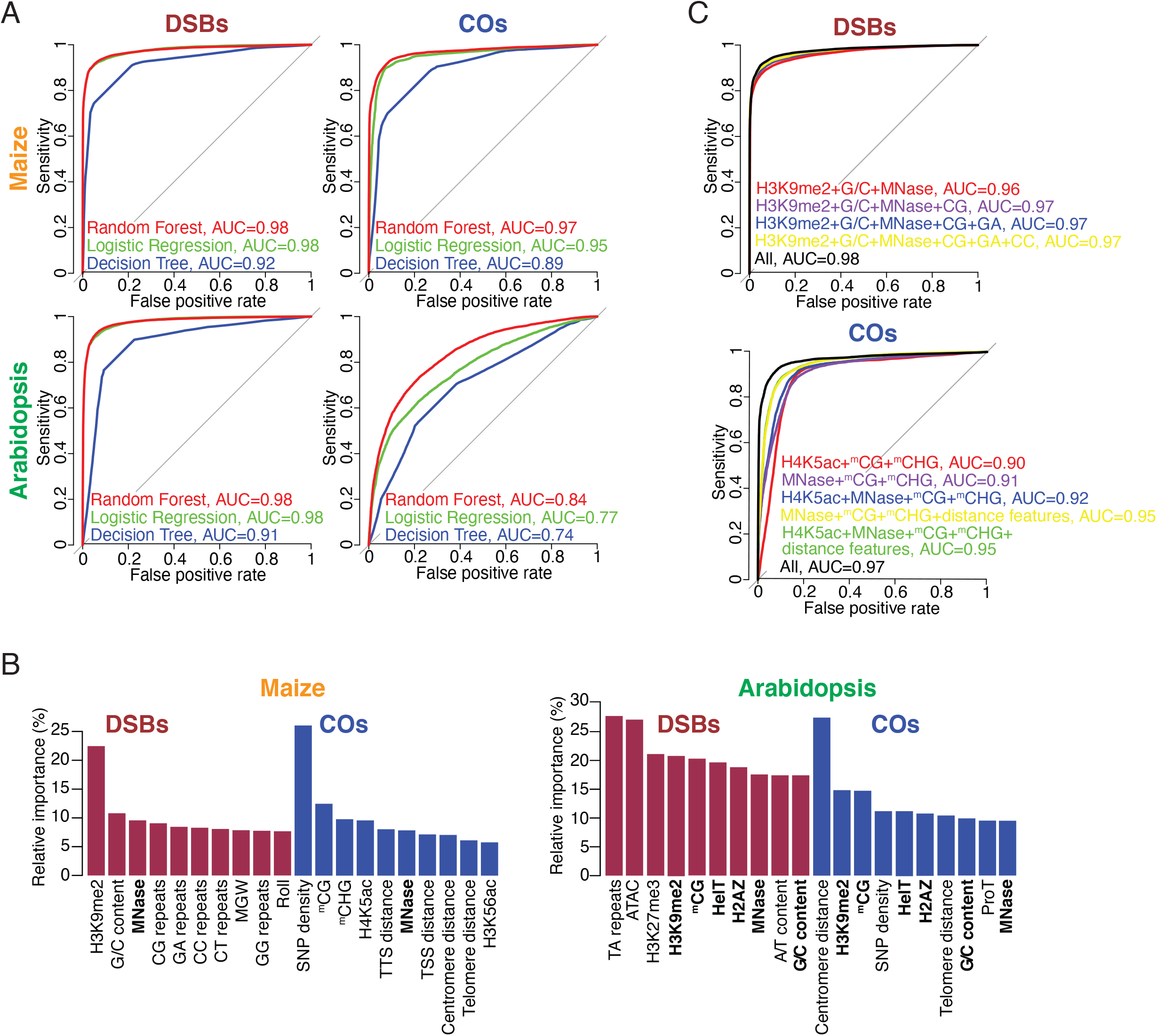
Using Machine Learning (ML) to identify features defining DSB and CO sites. **(A) Performance of three ML algorithms in identifying DSB and CO sites in maize and Arabidopsis**. Algorithm performance is measured by the Area Under Curve (AUC), which reflects the relationship between the algorithm’s detection ability (sensitivity) and the rate of false positives. AUC equal to 1.0 indicates perfect algorithm performance. **(B) Comparison of relative importance of ten features with the largest influence on defining the locations of DSBs and COs in maize and Arabidopsis**. Bold lettering identifies features that are among the ten most important for DSB and CO sites in each species. Note that we could not examine the impact of inter-parental sequence polymorphism on DSB formation, as the DSB datasets in the two species were obtained using homozygous strains^1,17^. **(C) Performance of the Random Forest algorithm in identifying DSB and CO in maize using a subset of genome and chromatin features**. Algorithm performance is measured by the Area Under Curve (AUC), which reflects the relationship between the algorithm’s detection sensitivity and the rate of false positives. AUC equal to 1.0 indicates perfect algorithm performance. G/C = G/C content; MNase = MNase sensitivity; CG, GA, and CC = frequencies of CG, GA, and CC dinucleotide repeats, respectively. Distance features are distance to the nearest transcription start and termination sites, telomere and centromere.

Of the ten genomic characteristics with most impact on CO site location, five were the same in maize and Arabidopsis: inter-parental DNA sequence polymorphism, DNA methylation at CG sites, nucleosome occupancy, and distance to centromere and telomere (Figure 1B, Supplemental Figure 4). However, only three of the ten most impactful DSB site characteristics, H3K9 di-methylation (H3K9me2), G/C content, and nucleosome occupancy, were shared by the two species. This difference suggests that CO site characteristics exhibit stronger evolutionary conservation than DSB site features. It should be noted, however, that different approaches were used for DSB detection in maize *versus* Arabidopsis^1,17^, which might have resulted in a lower between-species similarity of DSB site characteristics.

Interestingly, the ten features with strongest effects on DSB sites were not exactly the same as the ten features with strongest effects on CO sites, suggesting that distinct genome environments favor DSB formation *versus* their resolution into COs. In Arabidopsis, six of the ten features were shared by DSB and CO sites (Figure 1B). In maize, only one characteristics, nucleosome occupancy, was shared. H3K9me2, which had the strongest impact on DSB presence in maize of all features examined, was not among the ten most influential maize CO site features (Figure 1B). In contrast to yeast and mammals^8,21,22^, H3K4me3 was not a good predictor of either DSB or CO sites in our analyses, suggesting that plants may have a different hotspot recognition mechanism than yeast and mammals.

In maize, where the ML algorithms performed with more accuracy, we sought to determine the smallest number of features sufficient to reliably identify recombination sites. The combination of H3K9me2, nucleosome occupancy, and GC dinucleotide frequency allowed DSB detection with AUC of 0.96 (Figure 1C). In CO detection, the accuracy of 0.9 was achieved by the combination of DNA methylation at CG and CHG sites (where H is A, C, or T), together with either nucleosome occupancy or H4K5ac (Figure 1C). H4K5ac, which is known to facilitate formation of nucleosome-free regions in gene promoters^23^, was equivalent in its influence to nucleosome occupancy, and these two features exhibited strong correlation (Supplemental Figure 1). These results imply that few features are involved in recombination site designation and most features associated with recombination sites are redundant.

Using the minimal feature combinations allowed us to delineate genome regions capable of harboring DSBs and COs. We identified in the maize genome 154,520 candidate DSB sites and 64,301 candidate CO sites. Combined, these sites represent the recombination space, i.e. the fraction of the genome where meiotic recombination takes place. DSB space constitutes ∼ 13% of the maize genome while CO space is ∼ 5% of the genome. Interestingly, we found that the recombination space exhibits chromatin features that are very distinct from those of the genome at large. CG and CHG DNA methylation is normally strongly correlated with elevated levels of heterochromatin mark H3K9me2^24^. However, maize and Arabidopsis CO sites exhibit elevated H3K9me2 despite low DNA methylation (Figure 2). Furthermore, at DSB sites in maize, the level of H3K9me2 is low despite substantial levels of CG and CHG methylation (Figure 2).

**Figure 2.**
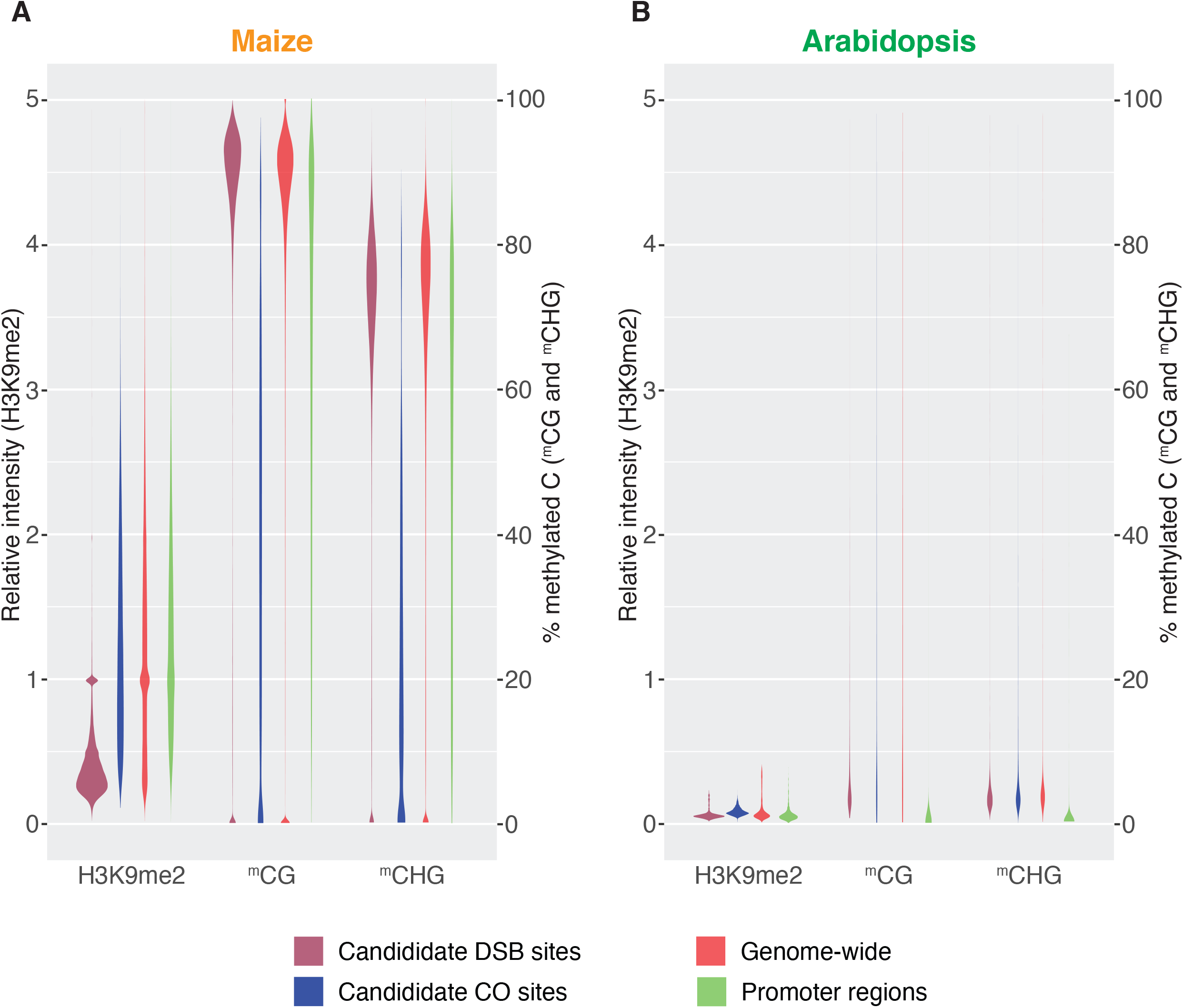
Comparison of H3K9me2 and DNA methylation at CG and CHG sites at candidate DSB and CO sites, 1000 randomly selected genome sites, and 1000 randomly selected promoter regions in maize (A) and Arabidopsis (B). All sites are 2 kb in size.

Over 80% of the candidate CO sites were located within 20 kb from a candidate DSB site and ∼70% were within 10 kb. However, only ∼ 30% of candidate CO sites actually overlapped candidate DSB sites. The median distance between a candidate CO site and an adjacent candidate DSB site was ∼ 4.0 kb. This shift may represent dHJ migration from its initial location at the DSB site to the place of its resolution. dHJ migration is a known feature of meiotic CO formation in yeast and mammals^25^ but has not yet been described in plants. Furthermore, we found that the fraction of CO sites that overlap DSB sites had lower levels of ^m^CG and H3K9me2 compared to COs located further away from DSB sites (Supplemental Figure 5). These observations further indicate that DSB formation and their resolution into COs are favored by distinct chromatin environments.

Studies in plants have shown that increased local DNA sequence polymorphism between parental chromosomes results in higher CO rates at specific loci^26-28^. There is, however, also evidence of DNA polymorphisms exerting anti-recombination effects^29^. Given the availability genome-wide CO data, we revisited this issue using, in addition to the maize and Arabidopsis datasets, CO maps of *indica* and *japonica* rice^30^. In all species, we found a clear relationship between SNP density and CO rates. Interestingly, there is a sequence polymorphism optimum at which the majority of COs are present, with fewer COs forming at sites with less and more polymorphism (Figure 3). Sequence polymorphisms in 2 kb as well as 100 kb windows around CO sites had similar effects. In the latter case, the optimum value of DNA sequence polymorphism roughly corresponds to the average SNP density in each species, accentuating the conclusion that CO formation favors sites with intermediate polymorphism levels. The mechanistic basis of the relationship between DNA sequence polymorphism and CO rates is not clear. CO formation could be associated with longer DNA overhangs produced by the single-end resection of meiotic DSBs^14^. Longer overhangs could tolerate more sequence mismatches^27^. Alternatively, CO distribution could be affected by patterns of chromatin remodeling factors, whose action is known to impact CO formation in yeast by controlling heteroduplex rejection when interparental DNA sequence polymorphism is excessive^31^.

**Figure 3.**
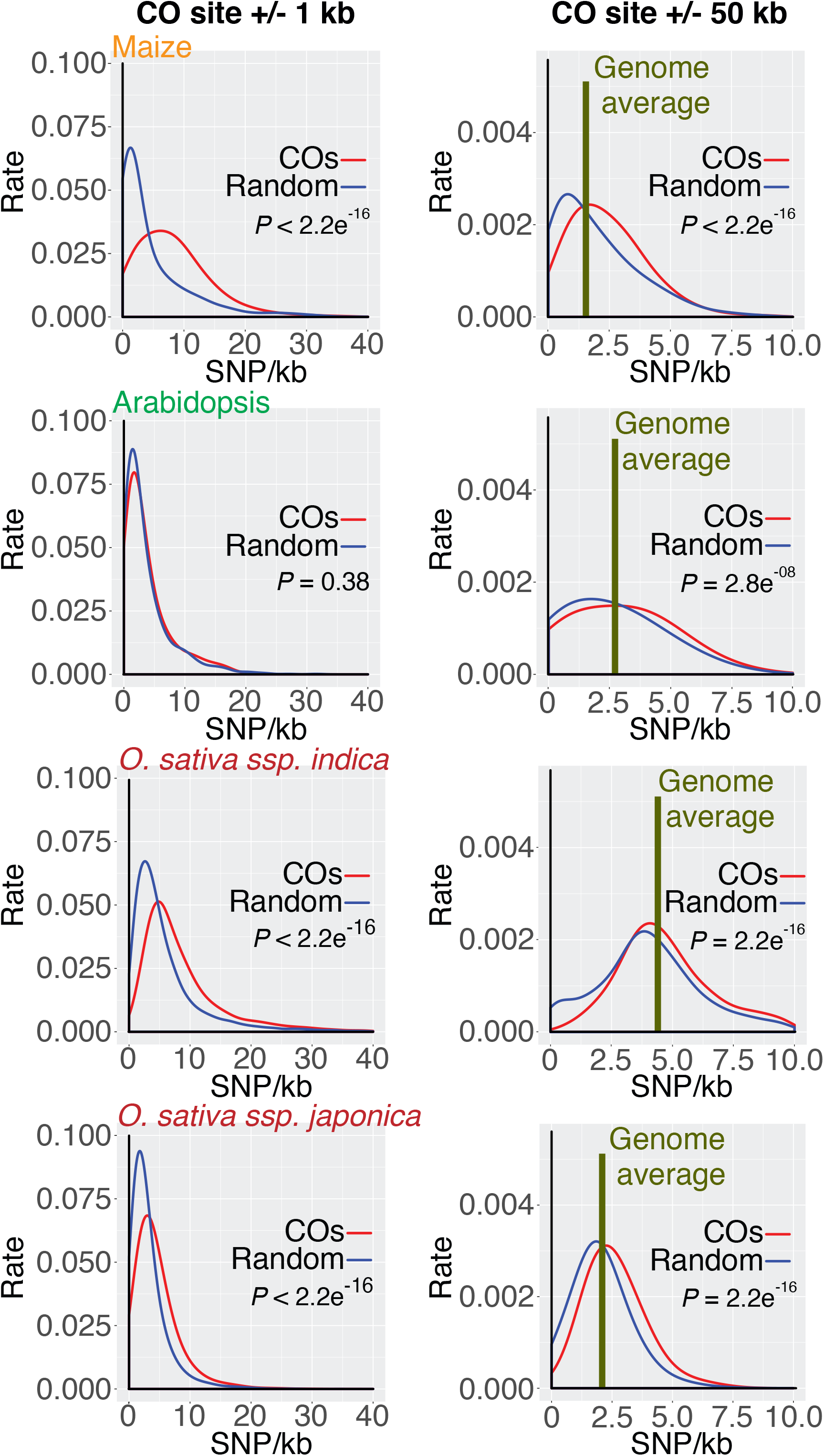
The relationship between CO frequency and DNA sequence polymorphism between parental chromosomes in 2 kb and 100 kb regions around CO sites. Probability values indicate the statistical significance of differences between CO distribution and the distribution of control samples generated by randomly selecting genome sites lacking COs. The number of sites in the control samples was the same as the CO number.

In both maize and Arabidopsis, the density of SNPs between parental chromosomes was among the ten most important factors impacting CO patterns. However, nearly the same CO site identification power could be achieved using the minimal algorithm that excluded the SNP density and relied solely on chromatin features. This observation suggests that inter-parental sequence polymorphism has either limited overall impact on CO landscape or that its effect is closely correlated with chromatin features.

To assess whether factors controlling CO locations vary among maize germplasm, we used the ML algorithm trained on the B73 x Mo17 CO dataset to detect COs in the B73 x CML228 hybrid^3^ as well as two diverse recombinant inbred line populations^32,33^. We found that 87% of COs reported in the three populations overlapped with CO sites predicted by the B73 x Mo17-trained algorithm (Supplemental Figure 6). The B73 x Mo17-trained algorithm was also able to correctly identify 93% of COs in Arabidopsis, 91% of COs in the *indica* rice population, and 87% of COs in the *japonica* rice population (Supplemental Figure 6). These data indicate that CO space characteristics exhibit limited within- and among-species variation.

Given this finding, we examined whether the locations of CO sites were also conserved. To do this, we determined the fraction of candidate CO sites associated with maize genes that are syntenic, i.e. exhibit conservation among grasses^34^. We found that, indeed, candidate CO sites tend to be in the vicinity of syntenic rather than non-syntenic genes (Supplemental Figure 7A). This result is consistent with the previous finding that most empirically-detected COs in maize are close to syntenic genes^35^. Candidate CO sites near syntenic and non-syntenic genes show distinct chromatin features and SNP levels (Supplemental Figure 7B, Supplemental Table 3). Altogether, these analyses indicate that CO space characteristics as well as location exhibited evolutionary conservation, implying uniformity of hotspot recognition mechanisms across plants. We also tested the B73 x Mo17 CO-trained algorithm on a set of 675 mouse COs mapped with the resolution of 2 kb or less^36^. We were only able to correctly identify only 87 of these COs, further accentuating the differences in recombination mechanisms between plants and mammals.

Taken together, we developed an ML protocol that is highly accurate in identifying sites of meiotic recombination events in plants based on genome sequence, chromatin, and chromosome location features. The algorithm performs equally well in genomes of varying sizes and complexities, including the small and lightly methylated genome of Arabidopsis and the fairly large and highly-methylated genome of maize. Our results indicate that CO landscape in plants can be explained in nearly its entirety by the combination of DNA methylation and nucleosome occupancy patterns. Interestingly, DNA methylation and nucleosome occupancy are not strongly associated with recombination sites in animals and fungi^8^. Furthermore, recombination sites in plants exhibit unique genome characteristics not found in other parts of the genome, particularly with respect to the interaction between DNA methylation and H3K9me2. Detailed recombination landscape maps are useful in studies of genome evolution and plant breeding efforts. The remarkable conservation of CO site features among plants, regardless of their genome size and complexity, may enable generating such maps in species in which creating genetic mapping populations is not feasible.

## MATERIALS AND METHODS

### Datasets

The maize CO dataset contained 1,136 COs mapped in a B73 x Mo17 BC1 population with the resolution of less than 2 kb^3^. The Arabidopsis CO set contained 1,078 events mapped to within 2 kb in a Col x Ler F2 population^2^. To infer CO positions in the maize dataset, a Hidden Markov Model (HMM) algorithm was used to correct for structural polymorphism between the B73 and Mo17 genomes^3^.

The maize DSB dataset contained 3,126 hotspots identified in the genome of the B73 inbred using chromatin immunoprecipitation (ChIP) with an antibody against the DSB marker protein RAD51^17^. To reduce false positives, the distribution of RAD51 ChIP-seq reads was compared with multiple negative control samples, including input chromatin, pre-immune IgG, and leaf tissue chromatin. Additionally, we used a stringent statistical cutoff value with a false discovery rate of <0.01. The source of Arabidopsis DSB hotspots was sequencing of SPO11-bound oligonucleotides released in the process of DSB formation^1^. For consistency, we processed sequencing reads from the Arabidopsis DSB dataset through a pipeline similar to the one we used to map DSBs in maize. Raw reads were trimmed using fastx_clipper to remove adapter sequences and aligned to the TAIR10 reference genome using bowtie2^37^ using the --very- sensitive setting and allowing for at most 2 mismatches. Replicate libraries were highly similar to each other (r=0.96 - 0.97 using 10-kb non-overlapping tiles throughout the genome) and were combined for further analysis. Single-end reads from a Columbia genomic DNA library were used as an input control after trimming to the length of 50 bp. MACS2 was used to conduct the enrichment analysis with the following parameter settings: --nomodel, --keep-dup=1 and -q 0.01. To be consistent with the size of the maize DSB hotspot dataset, the strongest 3,000 peaks were selected as the hotspot dataset. To be consistent with the CO datasets, DSB hotspots in both species were delineated as 2 kb regions centered on hotspot middle points.

### Chromatin and genomic feature data

All chromatin and genomic feature data used are listed in Supplemental Table 1. All sequencing data were mapped to reference genomes and filtered using a mapping quality score of Q20. Strength of each feature was calculated by subtracting input from a normalized treatment value. For nucleosome occupancy mapping, as the micrococcal nuclease-digested DNA results in much smaller DNA fragments than those of ChIP-derived DNA, we only focused on 1-kb wide regions Bisulfite sequencing data from DNA methylation analyses were trimmed to remove adapters, filtered to remove poor-quality reads and mapped to reference genomes using Bismark^38^ with default setting. The percent methylated cytosines in the CG, CHG or CHH contexts was calculated as the proportion of 100* (methylated reads)/ (methylated reads + unmethylated reads). Dinucleotide frequencies were computed using a Perl script. Distance to transcript start sites (TSSs), transcript termination sites (TTSs), centromeres and telomeres were calculated using the closest command of bedtools. The A/T and C/G contents were calculated using the nuc command of bedtools. DNA sequence motif frequencies were computed based on meme fimo results, with *P* < 1 e^-5^. The DNA shape structure of minor groove width (MGW), propeller twist (ProT), roll (Roll) and helix twist (HelT) were computed using a custom R script. To remove the bias arising from varying ranges of values of different features, each feature was normalized by scaling the minimum and maximum values to 0 and 1, respectively. In addition, features representing distance were log2 transformed before normalization. Feature patterns between positive (recombination sites) and negative (recombination deserts) datasets were comparted for each record using the Wilcoxon signed rank test implemented as wilcox.test in R.

### Construction of ML algorithms

Three different machine learning (ML) algorithms: Random Forest, Logistic Regression, and Decision Tree were trained to distinguish between positive (recombination sites) and negative (recombination deserts) datasets. To create negative control datasets, consisting of recombination deserts, we used the longest 5% of stretches between adjacent recombination sites. Regions annotated as containing centromeres were excluded, as centromeres may possess unusual features that block recombination^18^. To avoid bias, the same combined length of 2-kb long DNA fragments was randomly selected from the desert regions as the combined DNA length of the recombination site datasets.

All algorithms were executed in the R environment using 10-fold cross-validation and repeated 10 times. Datasets were randomly partitioned into 10 equal parts, and each part was successively used for testing of ML algorithm performance that was trained using the other 9 groups. To assess the prediction accuracy, area under the curve (AUC) was computed. Identification was considered correct when a predicted recombination site fell within 2 kb (i.e. the distance equal to the 2 kb resolution interval) from an empirically determined CO site or DSB hotspot.

Feature importance was estimated using the Random Forest algorithm and computed automatically using the mean as the model was built. To learn the model, feature values based on recombination sites were labelled as 1. Values from recombination desert regions were labelled as zero. The sets of minimal non-redundant features were examined using a forward sequential feature selection method with 10-fold cross validation. The contribution directions of features were assessed with the LR model using the sign of the beta coefficient.

### Genome-wide predictions using ML classifiers

Genome-wide predictions were performed by examining non-overlapping sliding windows of 2 kb in length, which corresponded to the sizes of DSB and CO intervals in the training datasets. To reduce false positives, we considered an interval a candidate DSB or CO site if it had a vote probability value (i.e. the fraction of trees indicating presence of a DSB or CO site) in the upper 25th percentile of the values above the default cutoff of 0.5. The candidate DSB and CO sites exhibited positive correlations with empirically detected DSB hotspots and COs at the 1-Mb scale (r = 0.48 for DSBs and r = 0.67 for COs; *P* < 0.001 by permutation). At the 20-kb scale, the correlations were also positive, albeit weaker (r = 0.08 for DSBs and r = 0.21 for COs; *P* < 0.001), most likely because of the relatively low number of empirically identified recombination events.

### Cross-genotype CO prediction

To assess whether factors controlling CO locations vary within maize, we used the ML algorithm trained on the B73 x Mo17 CO dataset to detect COs in diverse maize germplasm. We conducted the analysis on a set of 195 COs mapped with the resolution of 2 kb or less in the B73 x CML228 hybrid^3^ and two recombinant inbred line (RIL) populations, US-NAM and CN-NAM, in which more COs were mapped but with lower resolution, US-NAM and CN-NAM. US-NAM was created by crossing 25 genetically diverse inbred lines to B73^32^. CN-NAM, which was generated by crossing 11 Chinese breeding lines to Huangzaosi^33^.

## ACKNOWLEDGEMENTS

This research was supported by a grant from U.S. National Science Foundation (IOS-1546792) to WPP and the United States - Israel Binational Agricultural Research and Development (BARD) Fund (US-4828-15) to WPP and AAL.

## FIGURE LEGENDS

**Supplemental Table 1.** Sources of data used in the study.

| Category | Maize | Arabidopsis |
| --- | --- | --- |
| DSBs | He, *et al.*<sup>1</sup> | Choi *et al.*<sup>2</sup> |
| COs | Kianian, *et al.*<sup>3</sup> | Shilo *et al.*<sup>4</sup> |
| Genome features | B73v3 | TAIR10 |
| ATAC-seq | PRJNA336549 | PRJNA394532 |
| MNase-seq | GSE84368 | E-MTAB-5042 |
| DNA methylation | PRJNA357305 | PRJNA312219 |
| H2A.Z | - | PRJNA427312 |
| H3K4me3 | PRJNA185817 | PRJNA408288 |
| H3K9me2 | Zhou, *et al.*, in preparation | GSE12383 |
| H3K27me3 | PRJNA327594 | PRJNA418657 |
| H3K56ac | PRJNA327596 | PRJNA408284 |
| H4K5ac | PRJNA305809 | - |

1. He, Y. *et al.* Genomic features shaping the landscape of meiotic double-strand-break hotspots in maize. *Proc Natl Acad Sci USA* **114**, 12231-12236 (2017). 2. Choi, K. *et al.* Nucleosomes and DNA methylation shape meiotic DSB frequency in *Arabidopsis thaliana* transposons and gene regulatory regions. *Genome Res* **28**, 532-546 (2018). 3. Kianian, P.M.A. *et al.* High-resolution crossover mapping reveals similarities and differences of male and female recombination in maize. *Nat Commun* **9**, 2370 (2018). 4. Shilo, S., Melamed-Bessudo, C., Dorone, Y., Barkai, N. & Levy, A.A. DNA crossover motifs associated with epigenetic modifications delineate open chromatin regions in Arabidopsis. *Plant Cell* **27**, 2427-36 (2015).

**Supplemental Table 2.** Comparison of genome and chromatin feature characteristics at recombination sites and recombination deserts in maize.

| **Features** | **Wilcoxon rank sum test** |
| --- | --- |
| H3K4me3 | 6.228264e<sup>-61</sup> |
| H3K9me2 | 9.030779e<sup>-23</sup> |
| H3K27me3 | 3.664388e<sup>-270</sup> |
| H4K5ac | 1.016836e<sup>-191</sup> |
| H3K56ac | 8.599284e<sup>-113</sup> |
| MNase sensitivity | 6.859886e<sup>-167</sup> |
| ATAC | 3.048306e<sup>-146</sup> |
| Exon presence | 1.402454e<sup>-126</sup> |
| Transposable element (TE) presence | 1.348901e<sup>-27</sup> |
| SNP density | 7.966914e<sup>-257</sup> |
| Distance to nearest transcription start site (TSS) | 4.392102e<sup>-196</sup> |
| Distance to nearest transcription termination site (TTS) | 4.173331e<sup>-196</sup> |
| Distance to centromere | 4.799666e<sup>-80</sup> |
| Distance to telomere | 4.452987e<sup>-57</sup> |
| G/C content | 3.596657e<sup>-24</sup> |
| A/T content | 1.121656e<sup>-18</sup> |
| mCG | 4.682465e<sup>-263</sup> |
| mCHG | 1.591194e<sup>-236</sup> |
| Presence of DNA motifs | 5.515867e<sup>-112</sup> |
| AA repeats | 1.93321e<sup>-20</sup> |
| AC repeats | 1.421046e<sup>-19</sup> |
| AG repeats | 6.9798e<sup>-11</sup> |
| AT repeats | 2.546086e<sup>-07</sup> |
| CA repeats | 0.098110 |
| CC repeats | 0.011943 |
| CG repeats | 9.069393e<sup>-56</sup> |
| CT repeats | 5.666278e<sup>-06</sup> |
| GA repeats | 1.815234e<sup>-18</sup> |
| GC repeats | 2.805048e<sup>-59</sup> |
| GG repeats | 0.059243 |
| GT repeats | 6.178703e<sup>-10</sup> |
| TA repeats | 0.502548 |
| TC repeats | 5.2253e<sup>-12</sup> |
| TG repeats | 0.286701 |
| TT repeats | 2.798407e<sup>-16</sup> |
| Roll | 4.09787e<sup>-122</sup> |
| ProT | 1.443532e<sup>-11</sup> |
| HelT | 6.087287e<sup>-14</sup> |
| MGW | 9.724034e<sup>-86</sup> |
| Gene presence | 3.433425e<sup>-121</sup> |
| Single exon gene presence | $6.32463e^{-38}$ |
| Presence of gene expressed in meiosis | $4.185151e^{-12}$ |

**Supplemental Table 3.** Comparison of chromatin feature and SNP levels at CO sites in syntenic *versus* non-syntenic regions of the maize genome.

| Features | Probability of syntenic regions being different from non-syntenic regions |  |
| --- | --- | --- |
|  | ML-identified CO sites | Empirically-identified COs (Kianian *et al.*, 2018) |
| mCG | 4.32e<sup>-06</sup> | 2.47e<sup>-6</sup> |
| mCHG | 2.20e<sup>-16</sup> | 1.14e<sup>-5</sup> |
| Nucleosome occupancy | 0.04195 | 0.028 |
| H4K5ac | 2.20e<sup>-16</sup> | 7.75e<sup>-8</sup> |
| SNPs | 2.20e<sup>-16</sup> | 0.068 |

**Supplemental Figure 1.**
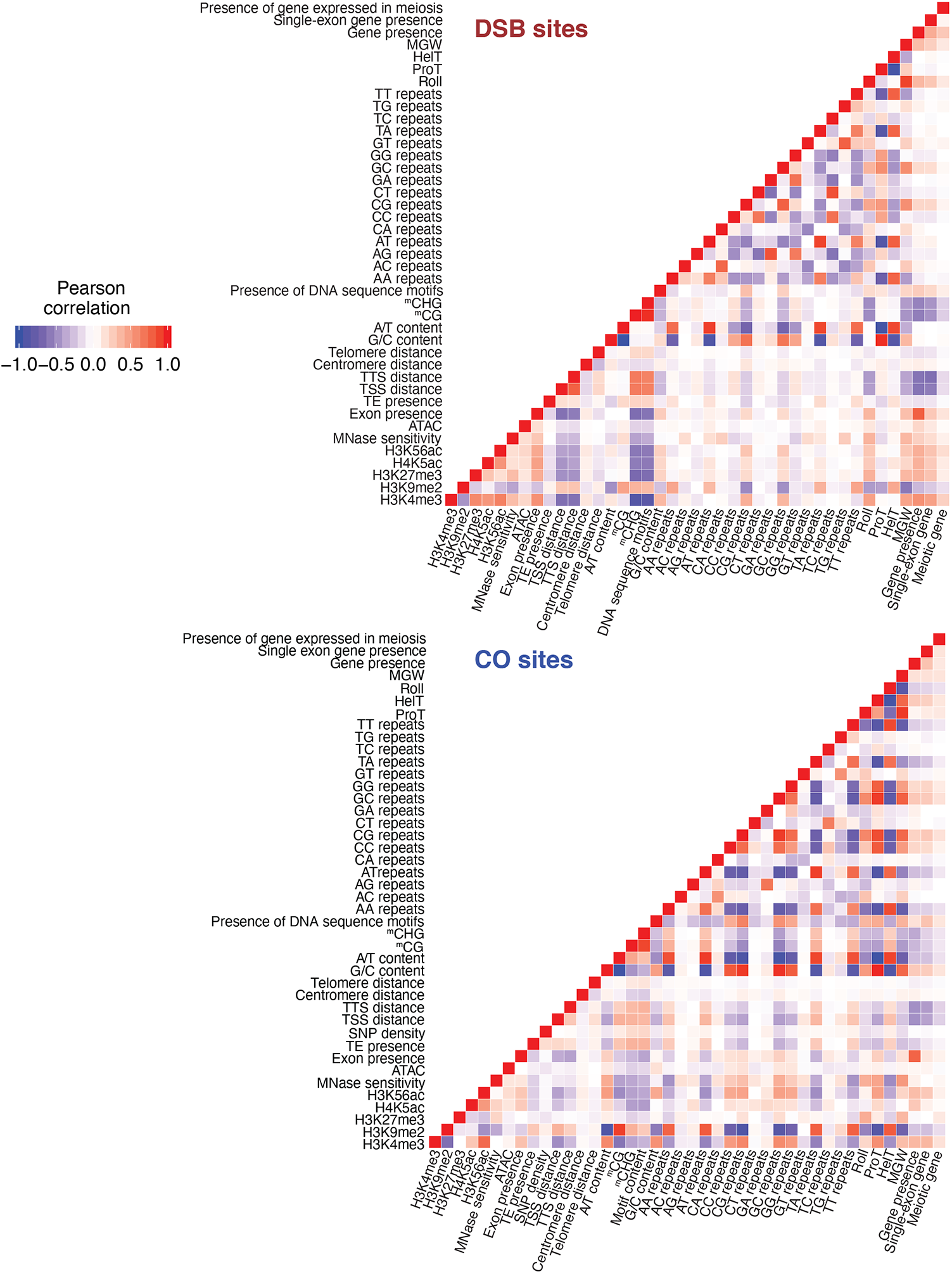
Pairwise correlations between genome and chromatin features in 2kb-long intervals centered at DSB hotspots and CO sites in maize. Pearson correlation coefficient values indicate that most recombination site features are only weakly correlated with each other. Nevertheless, some correlations between individual features can be detected, suggesting that these features redundantly represent the same chromosomal conditions.

**Supplemental Figure 2.**
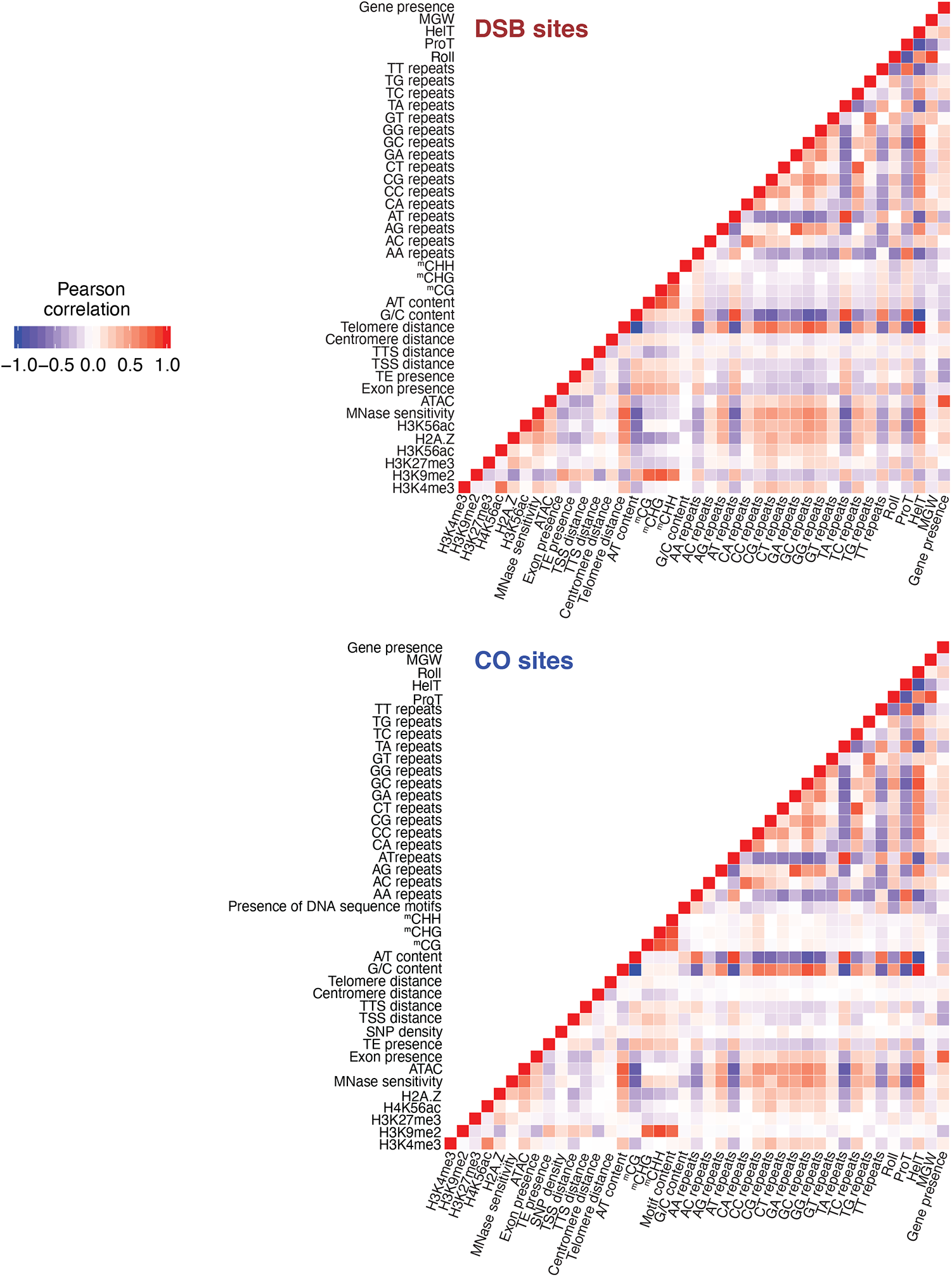
Pairwise correlations between genome and chromatin features in 2kb-long intervals centered at DSB hotspots and CO sites in Arabidopsis. Pearson correlation coefficient values indicate that most recombination site features are only weakly correlated with each other. Nevertheless, some correlations between individual features can be detected, suggesting that these features redundantly represent the same chromosomal conditions.

**Supplemental Figure 3.**
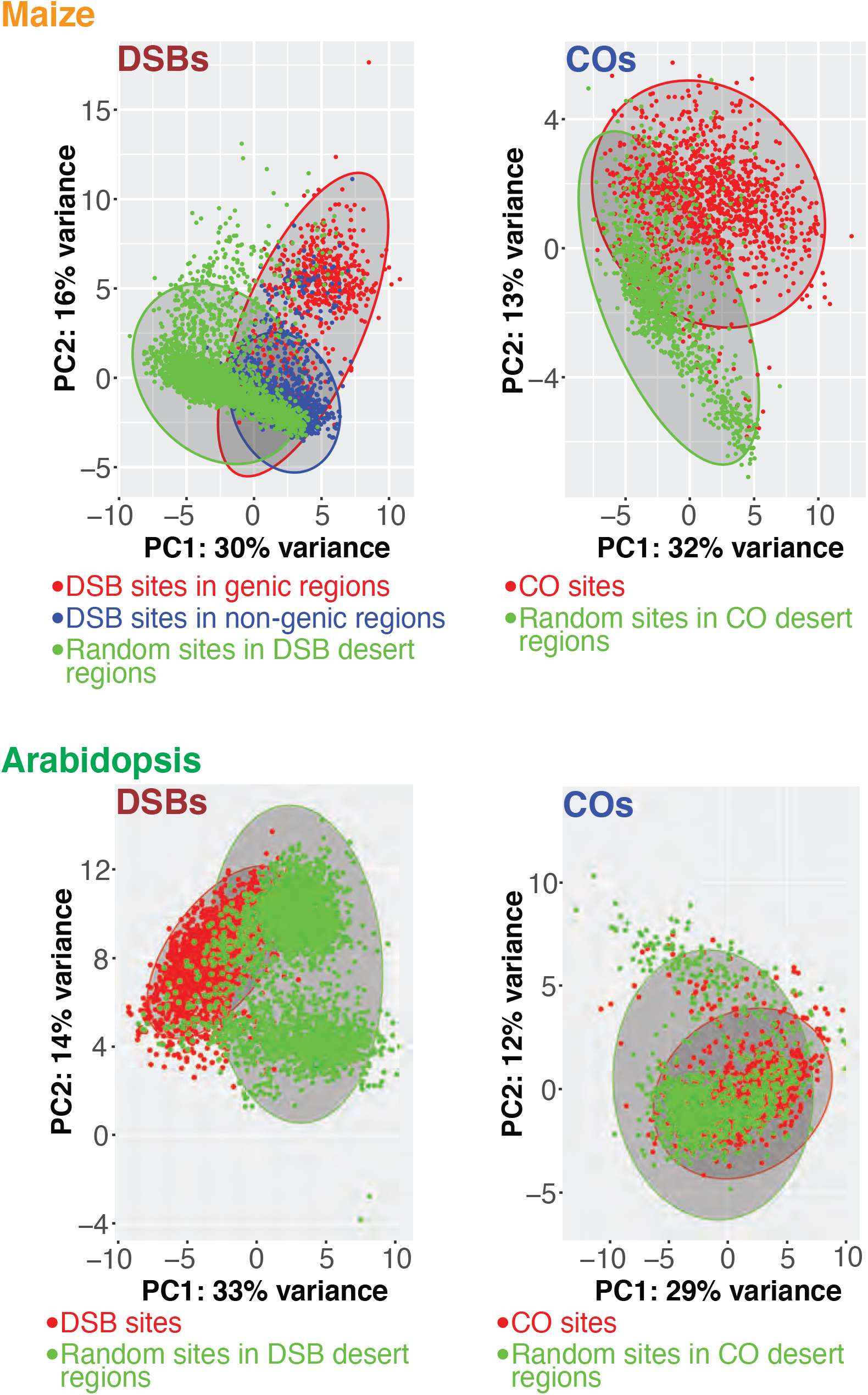
Principle component (PC) analysis of factors affecting locations of DSBs and COs. Features of DSB and CO sites in both maize and Arabidopsis exhibit strong separation from those of recombination desert regions. On the other hand, characteristics of DSB hotspots located in genic regions of maize largely overlap with those of DSB hotspots in repetitive DNA^1^, indicating that the overall DSB hotspot environments were similar, regardless of hotspot locations 1. He, Y. et al. Genomic features shaping the landscape of meiotic double-strandbreak hotspots in maize. *Proc Natl Acad Sci USA* 114, 12231-12236 (2017).

**Supplemental Figure 4.**
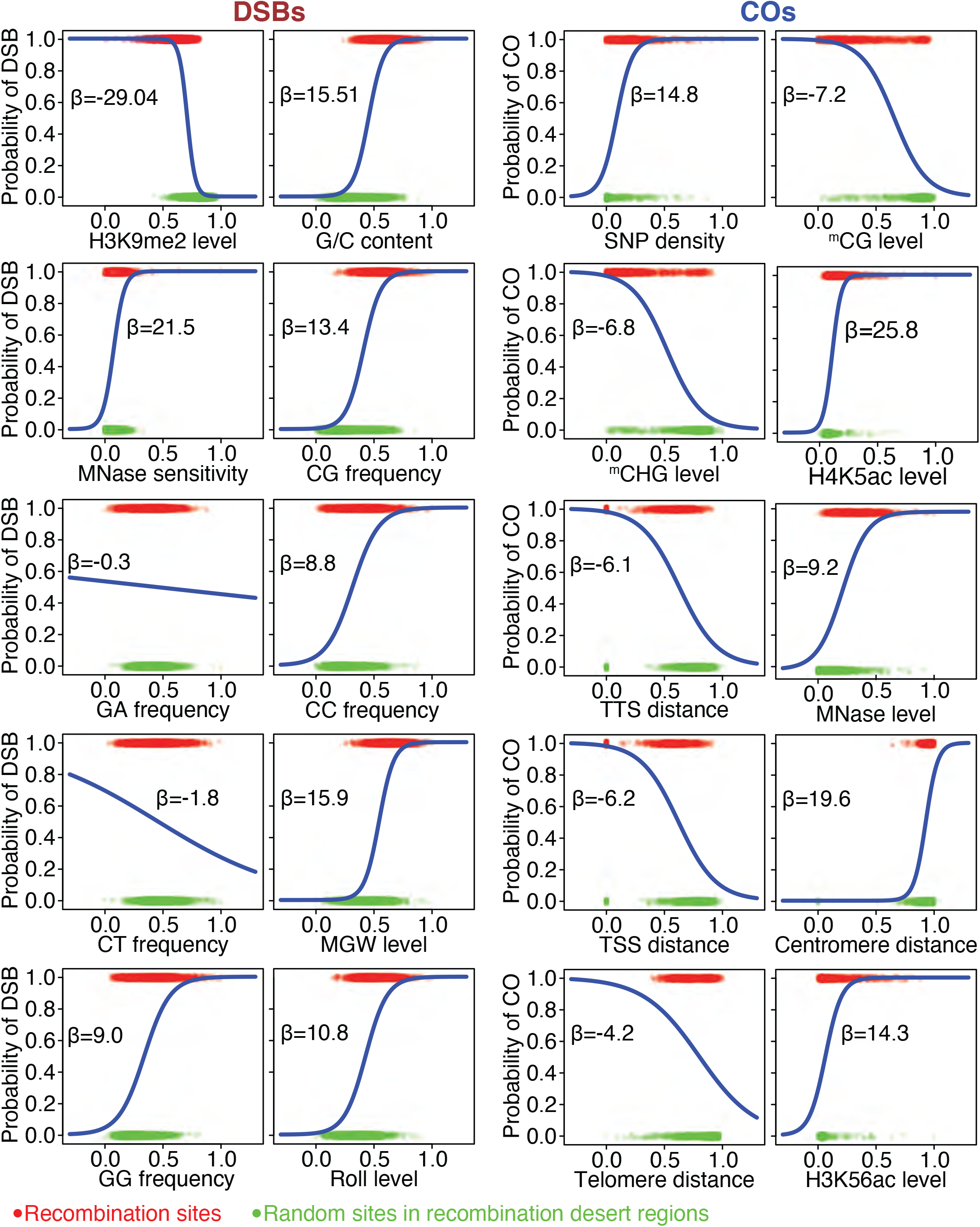
Effects of ten genome and chromatin features with the highest relative impact on DSB and CO locations in maize. X-axes represent normalized feature levels. Y-axes represent probabilities of DSB and CO site presence. As expected, features associated with active chromatin environment, such as low nucleosome occupancy, are positively correlated with the presence of recombination sites, whereas features associated with chromatin silencing, such as DNA methylation, are negatively correlated. Specific features show the same direction of influence on recombination sites for both DSBs and COs.

**Supporting Figure 5.**
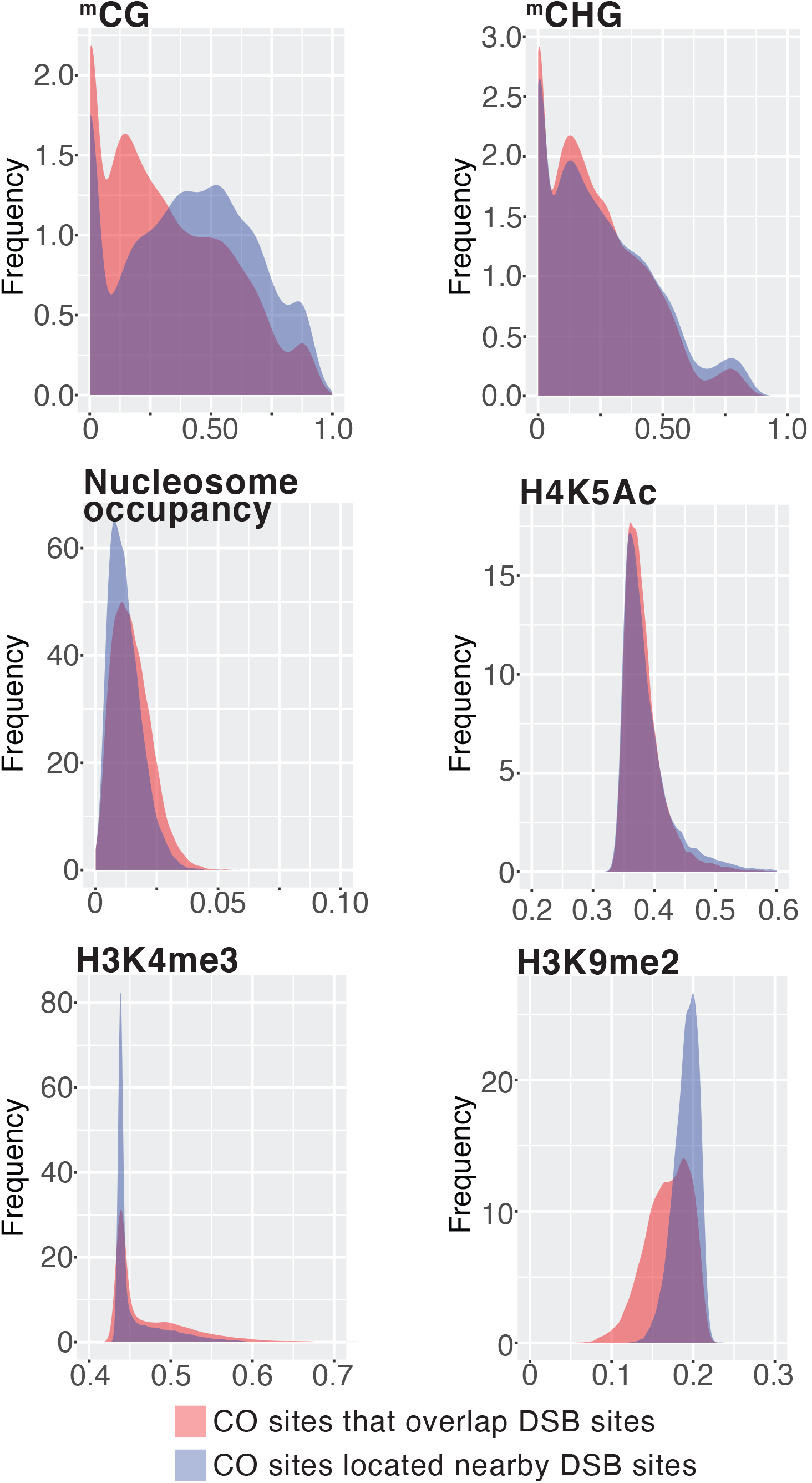
Comparison of chromatin features at candidate CO sites overlapping candidate DSB sites and CO sites located away from candidate DSB sites. X-axes represent normalized feature values. Y-axes represent frequency of sites exhibiting specific feature values.

**Supplemental Figure 6.**
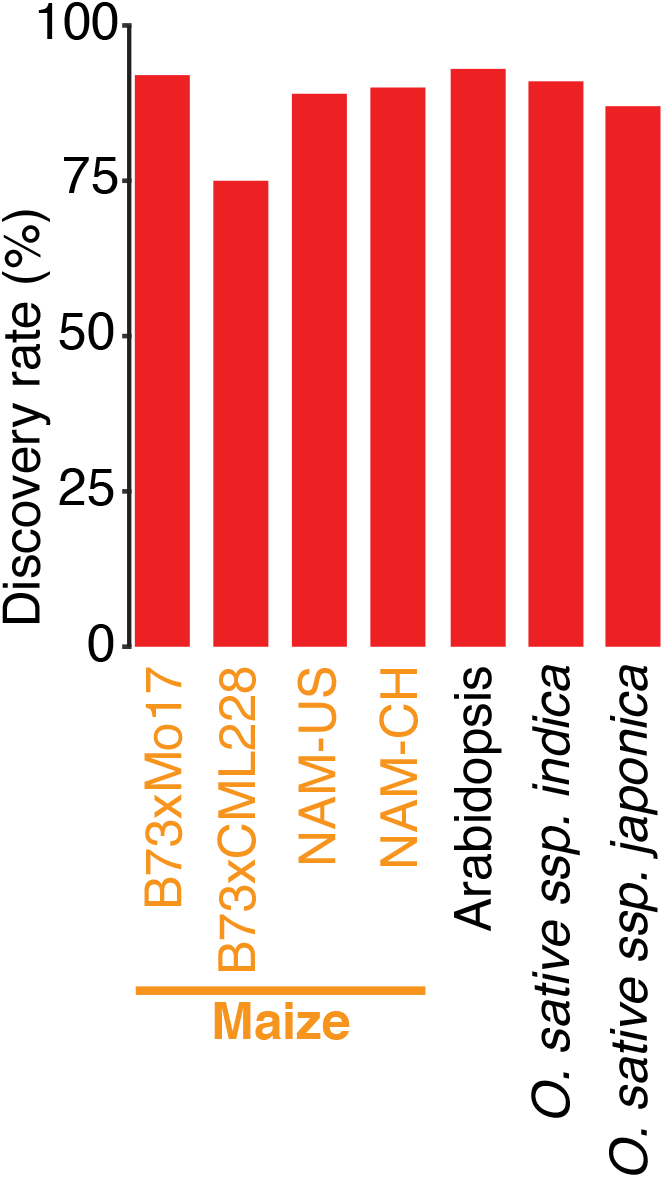
Comparison of CO site discovery rates in various maize populations and various species using the Random Forest algorithm trained on the B73 x Mo17 dataset.

**Supplemental Figure 7.**
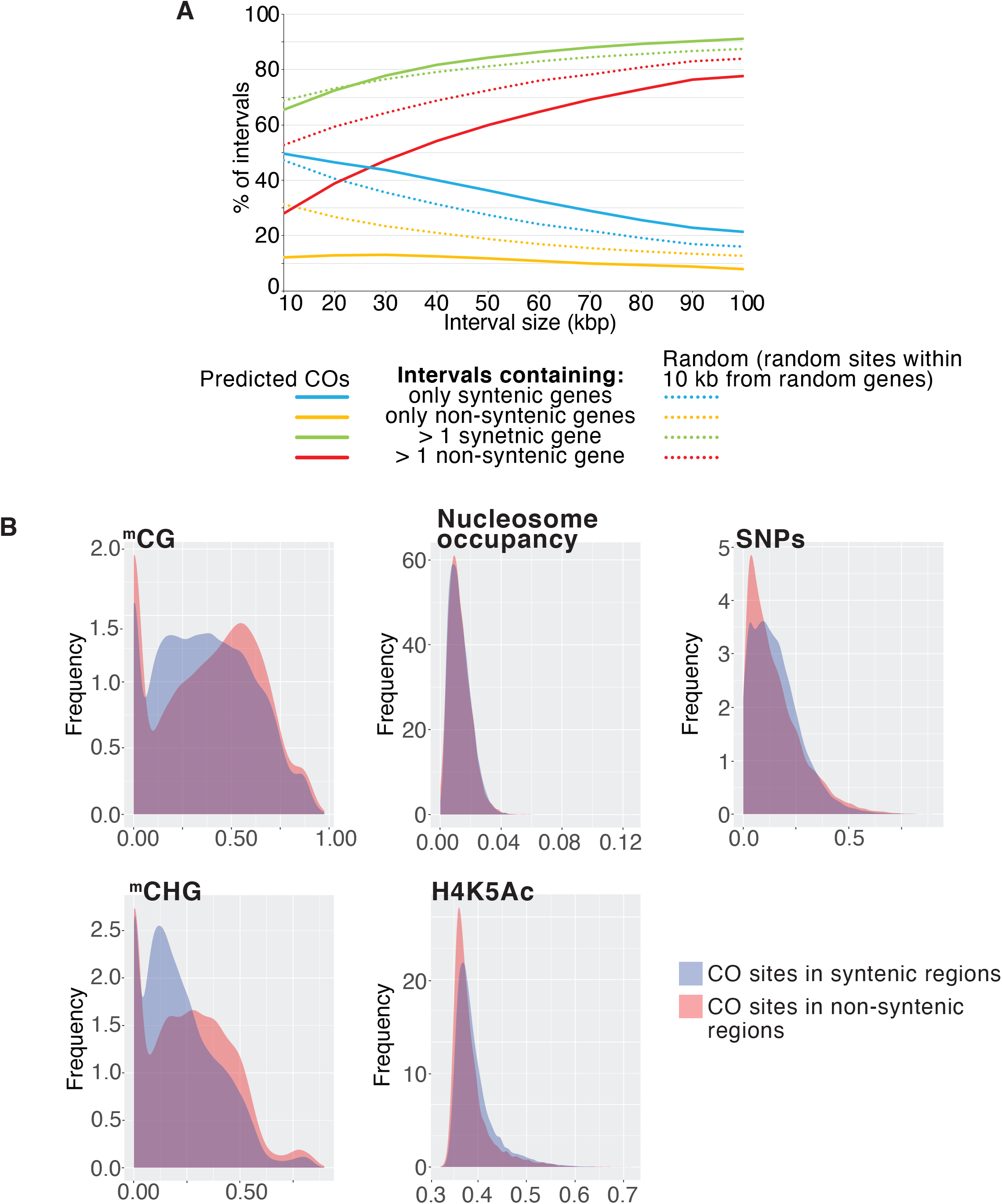
Candidate CO sites in syntenic and non-synetnic regions of the maize genome. (A) Proximity of predicted CO sites to syntenic and non-synetnic maize genes. (B) Comparison of chromatin features at candidate CO sites located in syntenic and non-syntenic regions of the maize genome. X-axes represent normalized feature values. Y-axes represent frequency of sites exhibiting given feature values.

